# Structural analysis of full-length SARS-CoV-2 spike protein from an advanced vaccine candidate

**DOI:** 10.1101/2020.08.06.234674

**Authors:** Sandhya Bangaru, Gabriel Ozorowski, Hannah L. Turner, Aleksandar Antanasijevic, Deli Huang, Xiaoning Wang, Jonathan L. Torres, Jolene K. Diedrich, Jing-Hui Tian, Alyse D. Portnoff, Nita Patel, Michael J. Massare, John R. Yates, David Nemazee, James C. Paulson, Greg Glenn, Gale Smith, Andrew B. Ward

## Abstract

Vaccine efforts against the severe acute respiratory syndrome coronavirus 2 (SARS-CoV-2) responsible for the current COVID-19 pandemic are focused on SARS-CoV-2 spike glycoprotein, the primary target for neutralizing antibodies. Here, we performed cryo-EM and site-specific glycan analysis of one of the leading subunit vaccine candidates from Novavax based on a full-length spike protein formulated in polysorbate 80 (PS 80) detergent. Our studies reveal a stable prefusion conformation of the spike immunogen with slight differences in the S1 subunit compared to published spike ectodomain structures. Interestingly, we also observed novel interactions between the spike trimers allowing formation of higher order spike complexes. This study confirms the structural integrity of the full-length spike protein immunogen and provides a basis for interpreting immune responses to this multivalent nanoparticle immunogen.

---

Severe acute respiratory syndrome coronavirus (SARS-CoV) caused a global outbreak from 2002-2003 causing severe pneumonia and killing almost 900 people (*1*). SARS-CoV-2, belongs to the same lineage of the β-CoV genus as SARS-CoV, and recently emerged in China, spreading rapidly and infecting more than 18 million people worldwide with cases continuing to rise each day (*2*). Given the global increase in population density, urbanization, and mobility, and the uncertain future behavior of the virus, vaccination is a critical tool for the response to this pandemic. The SARS-CoV-2 spike (S) trimeric glycoprotein is a focus of coronavirus vaccine development since it is a major component of the virus envelope, essential for receptor binding and virus entry, and a major target of host immune defense (*3, 4*). There are several currently ongoing efforts to make spike-based vaccines using different strategies (*4–6*).

The CoV S protein is synthesized as an inactive precursor (S0) that gets proteolytically cleaved into S1 and S2 subunits which remain non-covalently linked to form functional prefusion trimers (*7*). Like other type 1 fusion proteins, the SARS-CoV-2 S prefusion trimer is metastable and undergoes large-scale structural rearrangement from a prefusion to a thermostable post fusion conformation upon S-protein receptor binding and cleavage (*8, 9*). Rearrangement exposes the hydrophobic fusion peptide (FP) allowing insertion into the host cell membrane, facilitating virus/host cell membrane alignment, fusion, and virus entry. Notably, SARS-CoV-2 S has a 4 amino acid insertion (PRRA) in the S1/S2 cleavage site compared to SARS-CoV spike resulting in a polybasic RRAR furin-like cleavage motif that enhances infection of lung cells (*10, 11*). While the S2 subunit is relatively more conserved across the β-CoV genus, the S1 subunit comprising the receptor binding domain (RBD) is immunodominant and much less conserved (*12*). The FP, two heptad repeats (HR1 and HR2), transmembrane (TM) domain, and cytoplasmic tail (CT) are located in the S2 subdomain that encompasses the fusion machinery. The S1 subunit of SARS-CoV-2 S folds into 4 distinct domains; the N-terminal (NTD), the C-terminal domain (CTD) containing the RBD and two subdomains, SD1 and SD2. While some human CoVs (HCoV), including OC43, exclusively use NTD-sialic acid interactions as their receptor engagement, others like Middle East Respiratory Syndrome (MERS) CoV that use the CTD-RBD for primary receptor binding have also been reported to bind sialic acid receptors via their NTD to aid initial attachment to the host cells (*13–15*). Although SARS-CoV-2 primarily interacts with its receptor ACE2 through the CTD-RBD, there is currently no evidence indicating possible interactions between the NTD and sialoglycans (*16, 17*).

The structure of the stabilized SARS-CoV-2 spike ectodomain has been solved in its prefusion conformation and exhibits a high resemblance to SARS-CoV spike (*17–19*). In this report, we describe the atomic structure of a leading SARS-CoV-2 S vaccine candidate based on a full-length S gene with furin cleavage-resistant mutations in the S1/S2 cleavage site and the presence or absence of 2-proline amino acid substitutions at the apex of the central helix. Our studies reveal an overall shift in conformation of the S1 subunit compared to the previously published structures (*17–19*). Interestingly, we also observed direct interactions between adjacent spike trimers; the flexible loop between residues 615-635 in the SD2 from each trimer extending and engaging a binding pocket on the NTD of the adjacent trimers resulting in higher order spike multimers. Further, site-specific glycan analysis revealed the glycan occupancy as well as varying levels of glycan processing at the 22 N-glycosylation sequons present in the spike monomer. Thus, our studies provide in-depth structural analysis of the Novavax full-length vaccine candidate, currently being tested in humans, that appropriately recapitulates the prefusion spike.

## Design and validation of SARS-CoV-2-3Q-2P full-length spike

The SARS-CoV-2-3Q-2P full-length spike vaccine candidate (3Q-2P-FL) was engineered from the full-length SARS-CoV-2 spike gene (residues 1-1273) including the transmembrane domain (TM) and the cytoplasmic tail (CT) (Fig1a). The construct was modified at the S1/S2 polybasic cleavage site from RRAR to QQAQ to render it protease resistant along with 2 proline substitutions at residues K986 and V987 in the S2 fusion machinery core for enhanced stability (Figure 1A). A second full-length construct (3Q-FL) containing the cleavage site mutations but without the 2 proline substitutions was also made in parallel for comparative purposes. The FL spikes expressed and purified from insect cells were then formulated in 0.01% (v/v) polysorbate 80 (PS 80) detergent. To characterize the structural integrity of 3Q-2P-FL immunogen, we performed negative stain electron microscopy (nsEM) of the FL spike constituted in PS 80 in the presence of Matrix-M adjuvant, recapitulating the vaccine formulation being tested in humans. Micrographs and 2D classes of the particles revealed trimeric spike proteins present as free trimers or as multi-trimer rosettes with their transmembrane domains enclosed in cylindrical micellar cores of PS 80 detergent (Figure 1B). Micrographs show these 3Q-2P-FL spike nanoparticles containing as many as 14 trimers. The Matrix-M adjuvant can also be seen in the micrographs and 2D classes as spherical cages of different sizes with only very limited interaction with spike particles.

**Figure 1.**
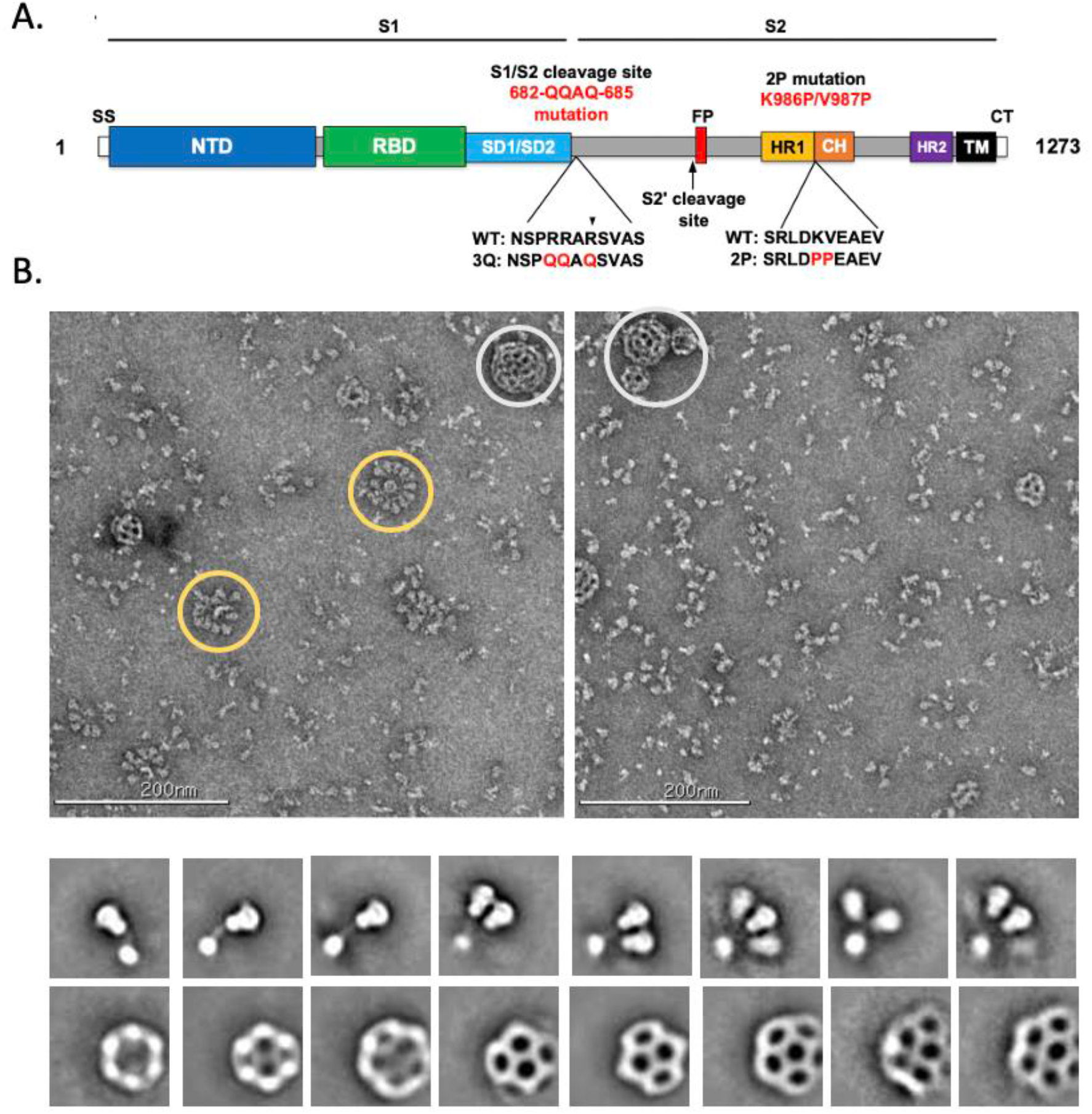
Evaluation of SARS-CoV-2 3Q-2P-FL spike glycoprotein. (**A**) Linear diagram of the sequence/structure elements of the full-length SARS-CoV-2 spike (S) protein showing the S1 and S2 ectodomain. Structural elements include a cleavable signal sequence (SS, white), N-terminal domain (NTD, blue), receptor binding domain (RBD, green), subdomains 1 and 2 (SD1/SD2, light blue), protease cleavage site 2’ (S2’, arrow), fusion peptide (FP, red), heptad repeat 1 (HR1, yellow), central helix (CH, brown), heptad repeat 2 (HR2, purple), transmembrane domain (TM, black) and cytoplasmic tail (CT, white). The native furin cleavage site was mutated (RRAR→QQAQ) to be protease resistant and stabilized by introducing two proline (2P) substitutions at positions K986P and V987P to produce SARS-CoV-2 3Q-2P-FL spike. (**B**) Representative negative stain EM images and 2D classes of SARS-CoV-2 3Q-2P-FL, formulated in polysorbate 80 detergent in the presence of Matrix-M adjuvant. In the raw micrograph, spike rosettes are circled in yellow and Matrix-M adjuvant cages are circled in white. 2D classes showing individual spikes, higher order spike nanoparticles and Matrix-M cages of different sizes.

## Cryo-EM of SARS-CoV-2-3Q-2P spike

To further evaluate the structural features of the 3Q-2P-FL immunogen, we performed single particle cryo-EM on the spike formulated in PS 80 detergent. The raw micrographs from cryo-EM revealed free trimers and trimer rosettes similar to those observed on the negative stain micrographs (Figure 2A). Initial 2D classifications revealed the presence of 2 distinct classes: free spike trimers and dimers of trimers (Figure 2A). Each class was independently subjected to additional classification and refinement.

**Figure 2.**
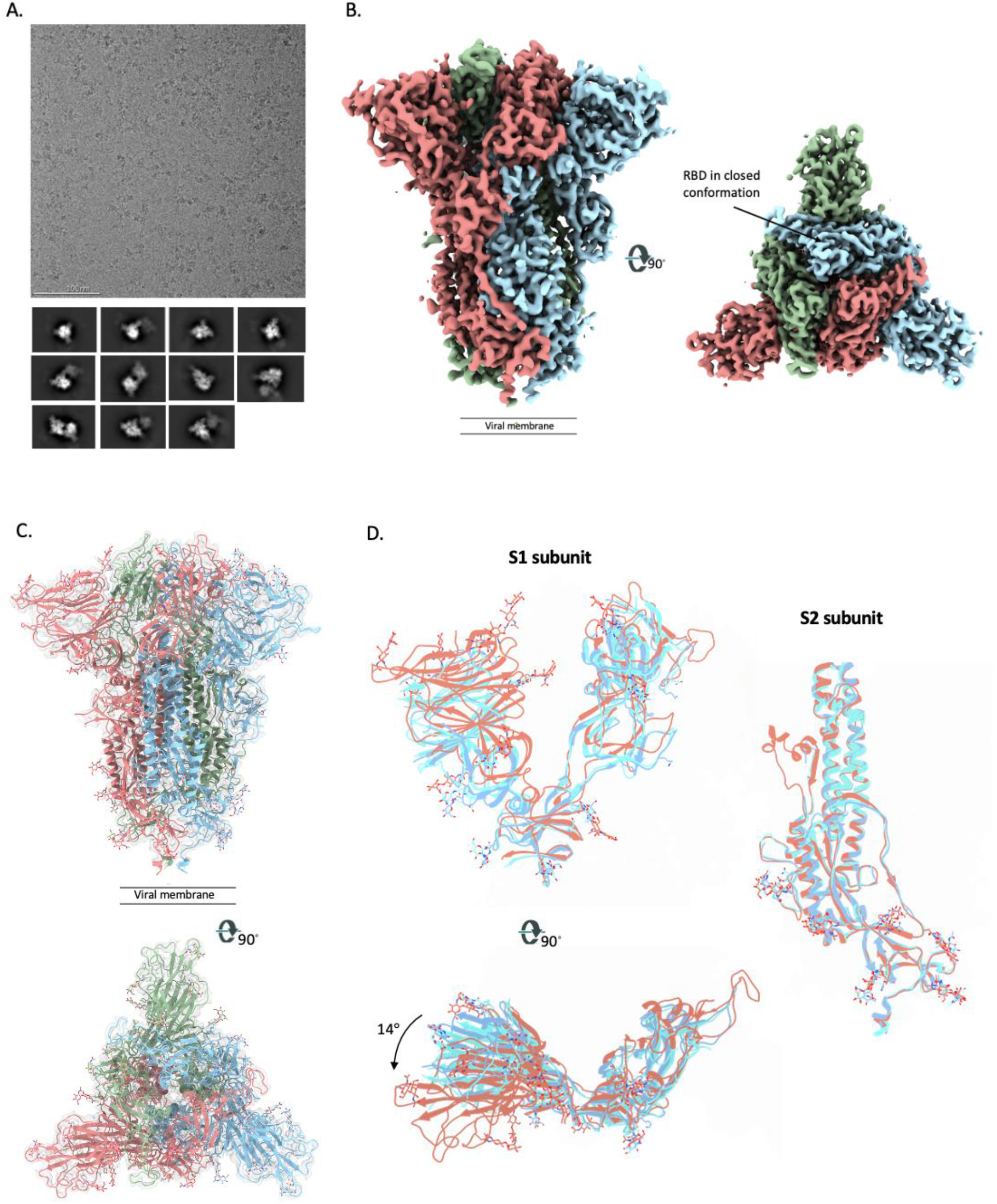
Cryo-EM analysis of SARS-CoV-2 3Q-2P-FL spikes. **(A)** Representative electron micrograph and 2D class averages of 3Q-2P-FL spikes showing free trimers and complexes of trimers **(B)** Side and top views of the B-factor-sharpened cryo-EM map of 3Q-2P-FL free trimers showing the spike in prefusion state, with the RBDs in closed conformation. The protomers are colored in blue, green and coral for clarity **(C)** Side and top view of the atomic model of free trimer represented as a ribbon diagram fit into the map density. The protomers are colored in blue, green and coral and the map is shown as a transparent gray density. **(D)** Comparison of 3Q-2P-FL spike with published structures (PDB IDs 6VXX and 6VSB) on a subunit level. PDB 6VXX is shown in cyan, PDB 6VSB shown in blue and 3Q-2P-FL spike in coral.

The three-fold symmetry (C3) reconstruction of the free spike trimer resulted in a map of 3.6 Å resolution while the asymmetric reconstruction (C1) was resolved to 3.8 Å resolution (Figure 2B and S1A, S1B). Previously published structures of soluble, stabilized SARS-CoV-2 spikes have revealed that RBDs exist in either a closed (RBD-down) or an open (RBD-up) conformation that can engage in ACE2 binding (*16–18*). In contrast, we observed that all three RBDs on the 3Q-2P-FL spike trimer were present in the closed conformation in the asymmetric reconstruction; the higher resolution C3 map was consequently used for model building (Figure 2B and S1C). Overall, the map was well resolved in both S1 and S2 subunits, particularly in the S1 NTD and CTD domains that were less resolved in previously published structures, thereby enabling us to model the full extent of these domains. Notably, the local resolution map calculated using cryoSPARC showed much of the spike trimer at substantially higher resolution than 3.6 Å (Figure S1D). The atomic model contains residues 14-1146 with breaks only in the flexible loop (619-631) and the cleavage site (678-688) (Figure 2C). Interestingly, superimposition of the coordinate models of 3Q-2P-FL spike with published spike structures (PDB Id: 6VXX and 6VSB) revealed substantial domain rearrangements in the S1 subunit of 3Q-2P-FL spike compared to the other models, whereas the structure of the S2 subunit was consistent with the published data (Figure 2D). The S1 NTD differed the most (~14° rotation counterclockwise relative to published models when viewing down towards the viral membrane) while the CTD and subdomains showed minor local rearrangements (Figure 2D). Notably, we also observed shifts in the placement of residues flanking the 615-635 loop compared to the published models. This region was modeled in one of the published structures (PDB Id: 6X6P) as a helix with residues flanking the helix positioned very differently from our model as shown by the corresponding placement of residues T632 and T618 (residues flanking the gap in 3Q-2P-FL model) (Figure 3A). However, upon closer inspection of the cryo-EM density (EMD-22078) corresponding to residues 621-640 of the PDB model 6X6P, there is insufficient density to support the helix conformation of this region (Figure S1E). The resulting displacement of residues in the 3Q-2P-FL structure enables inter-protomeric interactions by creating a salt-bridge between residues Asp 614 and Lys 854 (Figure 3B). This observation is particularly interesting given the increased prevalence of D614G mutation in the emerging SARS-CoV-2 strains and its potential role in viral transmission and pathogenesis (*20*).

**Figure 3.**
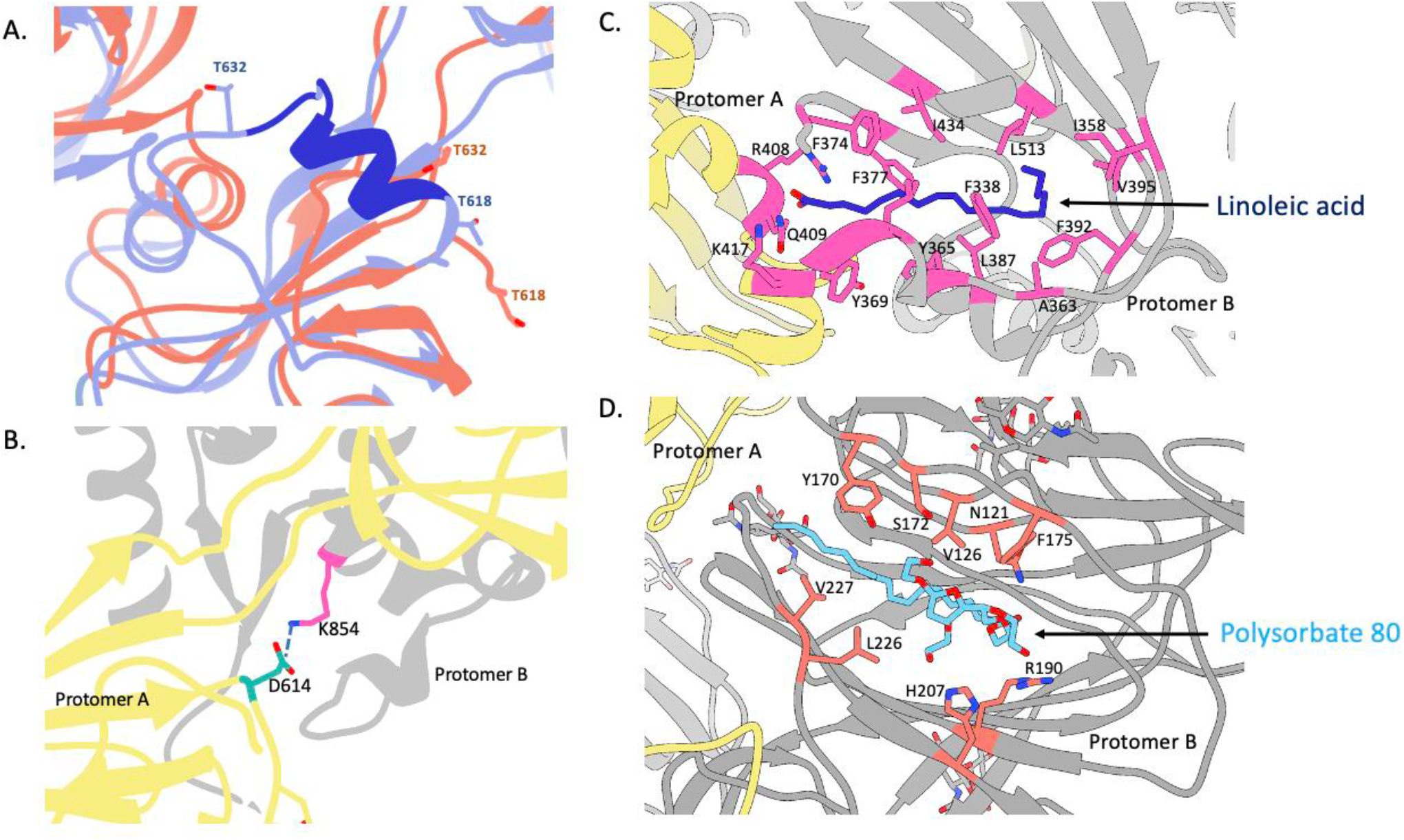
Structural features of the SARS-CoV-2 3Q-2P-FL spike trimer. **(A)**Comparison of the 615-635 loop between 3Q-2P-FL spike shown in coral and PDB 6X6P shown in blue. The residues that were built in 6X6P model but not in our model are shown in dark blue. Threonines at positions 618 and 632 flanking the gap in the 3Q-2P-FL trimer model are shown on both models to highlight their relative positions. **(B)** Interprotomeric salt-bridge interaction between D614 and K854 in 3Q-2P-FL spike trimer. **(C)** Linoleic acid (Dark blue) binding within a hydrophobic pocket of one RBD where the fatty acid head group reaches out to interact with the closed RBD of the adjacent protomer. The interacting residues are shown in pink **(D)**Polysorbate 80 detergent (blue) binding within the NTD with potential hydrogen bonding with R190 and H207. The interacting residues are shown in orange. Adjacent protomers are shown in yellow and gray in panels **(B), (C)** and **(D).**

During refinement of an atomic model into the EM density, we observed 2 additional densities in the S1 subunit that did not correspond to any peptide or glycans within the spike (Figure S2A). The first density was buried within a hydrophobic pocket of the CTD created by F338, F342, Y365, Y369, F374, F377, F392, F513 (Figure 3C and S2B). We had previously observed a non-protein density situated in the structure of porcine epidemic diarrhea virus (PEDV) that was identified to be palmitoleic acid (*21*).

This pocket in SARS-CoV-2 CTD corresponded with the structure of linoleic acid, a polyunsaturated fatty acid; the presence of this ligand was confirmed by mass spectrometry of 3Q-2P-FL spike (Figure S2B and S2C). The main chain carboxyl group of linoleic acid interacts with R408 and Q409 residues of RBD from the adjacent protomer thereby making interprotomer contacts (Figure 3C). The second unassigned density present in NTD was relatively larger and more surface exposed than the first density, surrounded by residues N121, Y170, S172, F175, R190, H207, V227 (Figure 3D and S2D). Analysis of the structural features of this density suggested that it may correspond to PS 80 detergent used to solubilize the membrane-bound trimers and stabilize them in solution. The aliphatic tail of PS 80 fit well into the hydrophobic pocket while the carbonyl and hydroxyl groups were well placed in proximity to residues R190 and H207 with potential for multiple hydrogen bonds between them (Figure 3D and S2D). Overall, the density is consistent with PS 80 detergent and, given its location, provides a possible explanation for the S1 shift seen in our FL trimer density compared to the published structures.

## Structures of dimer-of-trimers and trimer-of-trimers

Further classification of multimeric trimer particles yielded two separate classes; a dimer-of-trimers class that reconstructed to a final resolution of 4.5 Å with 2-fold symmetry and a trimer-of-trimers class that was resolved to 8.0 Å resolution (Figure 4A, 4B and S3A). The presence of the trimer-of-trimers class revealed that each spike trimer had the ability to interact with multiple trimers simultaneously. In both reconstructions, the interaction between each pair of trimers involved the SD2 of one protomer from each trimer engaging with the NTD of the adjacent trimer (Figure 4C). Consequently, each trimer pair is symmetrical along a 2-fold axis with trimer axes tilted to 44.5 degrees relative to each other. The atomic model of the dimer-of-trimer EM density revealed that the interaction was mainly coordinated by the 615-635 loop. Although most of the loop residues were too flexible to resolve in the free trimer density map, the inter-trimer interaction stabilized the loop so that it could be fully resolved (Figure 4D). The loop reaches into a pocket on the adjacent NTD, interacting with residues 621-PVAIHADQ-628 in the loop with NTD residues Q183, H146, Y248, L249, V70 and S71 (Figure 4D). We observed subtle changes in the NTD binding pocket in the loop-bound state compared to the free trimer model that allow better accommodation of the loop in the pocket. The residues Y145 and H146 in the binding pocket appear to switch positions in the loop-bound state resulting in a salt bridge interaction between H146 and D627 and potential stacking between W152 and H146 (Figure 4E). We also observed minor displacement of residues 68-75 and 248-250 surrounding the pocket. In addition to the main loop interaction resulting in higher-order oligomers, we also observed N282 glycans extending out towards the symmetry related chain in the adjacent trimer (Figure S3B). Parts of the glycans are stabilized and visible in the dimer-of-trimers map. Due to the close proximity, these glycans might form hydrogen bonds with the symmetry related chain but it is unclear if these glycan interactions aid in the stability of the dimer-of-trimers.

**Figure 4.**
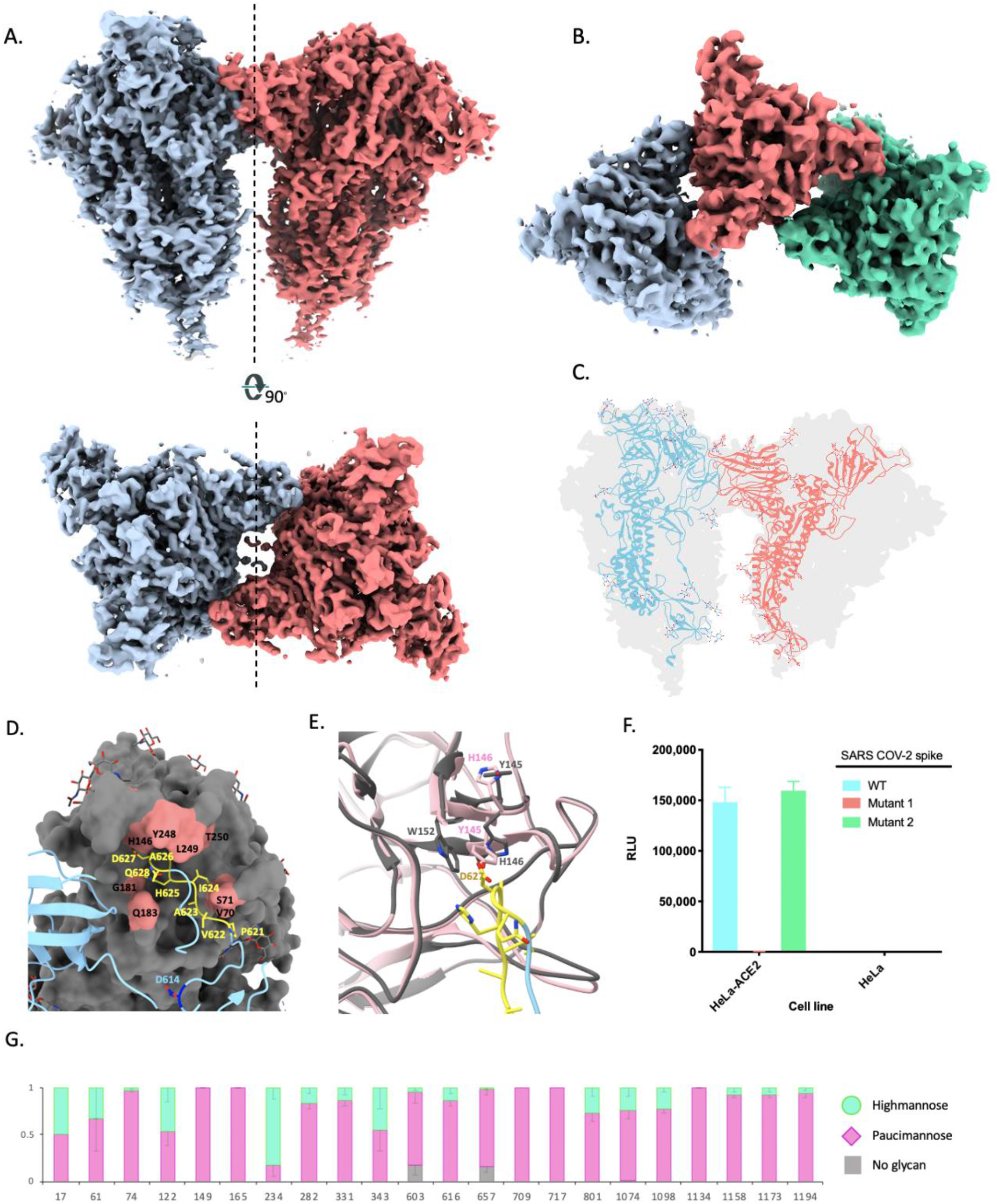
Trimer-trimer interactions and glycan analysis. **(A)**Side and top views of the sharpened cryo-EM map of 3Q-2P-FL dimers of spike trimers. Individual spike trimers are shown in blue and coral along a two-fold axis of symmetry (dotted line). **(B)** Top view of the B-factor-sharpened cryo-EM map of trimer-of-trimers complex with individual trimers colored in blue, coral and green. **(C)** Ribbon representation of a protomer from one trimer (blue) interacting with the protomer from the adjacent trimer (coral) docked into the dimers-of-trimers density **(D)** A close-up view of the interaction between the protomers of adjacent trimers. One protomer is shown as a ribbon diagram in blue while its binding partner is shown as surface in gray. Residues 621-PVAIHADQ-628 in the loop with potential interactions to the neighboring NTD are colored yellow and the residues in the NTD binding pocket are highlighted in coral. Residue D614 at the start of the loop is highlighted in dark blue. Glycosylation at residue 616 is not shown for clarity. **(E)** Changes occurring in the binding pocket in the bound state (gray) versus the free trimer (pink). Y145 and H146 switch positions to accommodate the loop better, also resulting in salt bridge formation between H146 and D627. It also results in stacking between W152 and H146 **(F)** Pseudoviruses expressing SARS-CoV-2 WT or mutant spikes (Mutant 1 has loop residues 621-PVAIHADQ-628 replaced with a glycine-serine linker and mutant 2 has residues 619-EVPV-622 mutated to 619-DVST-622) were used to infect HeLa or HeLa-ACE2 cells for 42 to 48 hours. Infection was measured by luciferase intensity (Relative Light Unit) in the lysed cells following infection. **(G)** Site-specific glycan analysis of 3Q-2P-FL spike protein expressed in SF9 insect cell line. Proportions shown for no occupancy, oligomannose and complex/paucimannose PNGS are the average and SEM of 3-32 unique peptides for each glycosite except for sites 17, 709 and 717 where only a single peptide was observed.

To investigate if the residues involved in trimer-trimer interactions are conserved across CoV strains belonging to lineage B of betacoronaviruses, we performed sequence alignment of residues in the loop and corresponding NTD binding pocket across representative strains (Figure S3C). While the loop residues 621-PVAIHADQ-628 are well conserved across strains, we observed varying levels of conservation in the binding pocket with SARS-CoV exhibited the most variability. SARS-CoV-2 has 7 additional residues at position 70-76 (including V70 and S71), 4 additional residues at position at 150-153 and 6 additional residues at 247-252 (including Y248 and L249) compared to SARS-CoV. These insertions along with the absence of H146 at the corresponding site on SARS-CoV make it unlikely for SARS-CoV to participate in these inter-trimeric interactions seen in SARS-CoV-2. While the residues in the NTD pocket were almost identical between SARS-CoV-2 and its closely related bat strain Bat-SL-RatG13, we observed some residue differences and 1-3 amino acid deletions in the loops comprising the NTD binding pocket of representative strains Bat-SL-CoVZC45, BetaCoV/pangolin/Guandong/1/2019 and BetaCoV/pangolin/Guangxi/P4L/2007 that could potentially impact binding to the loop (Figure S3C).

Due to the proximity of this binding pocket to the putative glycan binding site, we conducted structural comparisons of the NTD dimerization pocket with that of the sialic acid (SA) binding site on MERS spike (PDB ID: 6Q04) (*15*). Comparison to MERS showed that although the SA-binding pocket was in close proximity, it did not coincide with the dimerization pocket on NTD (Figure S3D). While it has not been identified if SARS-CoV-2 spike binds to glycans in a similar manner, computational and structural studies have proposed residues on SARS-CoV-2 spike involved in SA binding (*22, 23*). Structural comparison of this putative glycan binding site to the dimerization site revealed them situated adjacent to one another with residues in loop 70 contributing to both the binding pockets (Figure S3E).

To evaluate if the multimerization phenomenon observed in the full-length spike construct played a role in virus replication, we performed pseudovirus replication assays with SARS-CoV-2 wild-type (WT) spike and two mutant spikes. In mutant 1, the loop residues 621-PVAIHADQ-628 were replaced with a glycine-serine linker to completely knockout binding to NTD and in mutant 2, residues 619-EVPV-622 of SARS-CoV-2 were reverted to residues 619-DVST-622 of SARS-CoV-1. Pseudoviruses containing either WT or mutant spikes were generated in HEK293T cells and used to infect HeLa or HeLa-ACE2 cells. While the WT and mutant 2 exhibited similar levels of infection, we observed no detectable levels of infection for mutant 1 in which all contact residues on the loop were replaced by a GS linker (Figure 4F).

## Structural comparison to SARS-CoV-2-3Q-FL spike

To investigate if the absence of 2 stabilizing proline mutations impacted the spike stability and formation of higher order multimers, we performed cryo-EM studies of the SARS-CoV-2-3Q-FL (without 2P) protein formulated in PS 80 detergent. The raw micrographs and 2D classes revealed the presence of free trimers as well as trimer-trimer complexes as observed with 3Q-2P-FL, indicating that the proline stabilization is not necessary for the formation of these higher order complexes (Figure S4A). The 3D refinement of free trimers was refined to 4 Å resolution imposing C3 symmetry as we observed that the RBDs were present in closed conformation similar to 3Q-2P-FL (Figure S4B). Fitting the 3Q-2P-FL model into the 3Q-FL map revealed identical conformation of the spike protein further supporting that the presence of 2P in the full-length immunogen does not lead to any structural changes in the spike protein (Figure S4C).

## Glycan occupancy and glycan processing

Glycans on viral glycoproteins play a wide role in protein folding, stability, immune recognition and potentially in immune evasion. Site-specific glycosylation of the SARS-CoV-2 prefusion spike protein produced in SF9 insect cells was analyzed using our recently described mass spectrometry proteomics-based method, involving treatment with proteases followed by sequential treatment with the endoglycosidases (Endo H and PNGase F) to introduce mass signatures in peptides with N-linked sequons (Asn-X-Thr/Ser) to assess the extent of glycosylation and the degree of glycan processing from high mannose/hybrid type to complex type (*24*). Although the method was developed to assess the degree of processing of N-linked glycans in mammalian cells, it is also applicable for analyzing glycosylation of SF9 insect cells. The primary differences in glycan processing of N-linked glycans in SF9 insect cells are: 1) the production of truncated paucimannose glycans, and 2) the potential to introduce either one (α1,6) or two (α1,6/α1,3) fucose substitutions into the core GlcNAc attached to Asn. Although α1,3 fucose substitution is known to prevent cleavage by PNGase F (*25*), this is not a factor when analyzing glycosylation from SF9 cells since they contain α1,6-fucosylatransferase, which is found in mammalian cells, but only contain trace amounts α-1,3-fucosyltransferase activity, if any (*26*). The paucimannose glycans are highly processed like complex type glycans and not cleaved by EndoH, but are cleaved by PNGase F. Thus, for SF9 insect cell-produced glycoproteins, the use of endoglycosidases to introduce mass signatures is analogous to analysis of glycoproteins produced in mammalian cells, with EndoH removing high mannose/hybrid glycans leaving a GlcNAc-Asn (+203), followed by treatment with PNGase F in O_18_ water which removes the remaining paucimannose and complex type glycans and while converting Asn to Asp (+3), and the Asn of unoccupied sites remains unaltered (+0).

Our analysis detected glycosylation at all 22 potential N-linked glycan sequons present on SARS-CoV-2 spike (Figure 4G). Overall, there was high glycan occupancy of over >98%, with only two sites, 603 and 657, more than 5% unoccupied. Interestingly, we did not see clear glycan density at either 603 or 657 in the cryo-EM reconstruction of the 3Q-2P-FL spike. Most sites showed extensive glycan processing to complex/paucimannose type glycans, with only four sites that exhibit ≥40% oligomannose. The glycan analysis also confirmed the presence of glycans at sites 1158, 1173 and 1194 present in the membrane-proximal region of the spike not resolved by cryo-EM. The extensive site-specific glycan processing of the SARS-CoV-2 prefusion spike protein in SF9 insect cells seen here is similar to that recently reported for the spike protein produced in mammalian HEK293F cells (*27*).

## Discussion

The coronavirus disease (COVID-19) caused by SARS-CoV-2 poses a serious health threat and was declared a pandemic by the World Health Organization (WHO). In quick response to this rapidly evolving situation, several SARS-CoV-2 spike-based vaccine candidates are being developed and tested at various stages of clinical trials (*4-6*). In this study, we performed structural analysis of the Novavax SARS-CoV-2-3Q-2P full-length immunogen formulated in polysorbate 80 detergent. Our results show that PS 80 forms detergent micelles around the transmembrane domains of 1 or more spike proteins, enabling the formation of nanoparticle-like rosettes. These larger nanoparticles provide multivalent display of the spike immunogen with potential for better immunogen trafficking and B-cell activation as has been previously shown for nanoparticle-based immunogens (*28–30*). Consistent with this observation, analysis of safety and immunogenicity of the Novavax SARS-CoV-2-3Q-2P-FL immunogen in mice and baboons revealed strong B-and T-cell responses to the vaccine with no evidence of vaccine-associated enhanced respiratory disease (VAERD) (*31*).

Structural analysis of the 3Q-2P full-length spike immunogen revealed several important findings. The first being the stabilization and shift of the S1 subunit compared to published structures (*17, 18*). Although the cause of this shift is unclear, the presence of PS 80 wedged in the NTD potentially may stabilize the alternate conformation. Notably, another recent study observed differences in their NTD conformations as a function of pH (*32*). We also observed intertrimer interactions between SARS-CoV-2 full-length spike proteins for the first time. The 615-635 loop that is generally disordered in free trimers, engages the NTD of the adjacent trimer in a well-ordered conformation. The binding pocket on NTD is present adjacent to the putative glycan binding site and shows subtle differences in the apo and loop-bound conformations. The slight displacement of loop residues 68-75 and the reversed positions of Y145 and H146 in the bound state not only allow better accommodation of the loop in the pocket, but also enable the formation of salt bridge interactions between H146 and D627. Importantly, both these findings were seen in the full-length spike immunogen assembled into compact and dense nanoparticles, which may play a role in both the observed S1 shift as well as formation of higher order spike multimers. Cryo-electron tomographic reconstructions of intact SARS-CoV-2 virions showed a relatively dispersed distribution of spike protein trimers on the viral surface and no evidence of higher order aggregates (*33*). However, another study has shown that the D614G mutation present in close proximity to the dimerization loop results in a several fold increase of spike numbers on the viral surface, resulting in higher spike protein density and a more infectious virion (*20*). The greater density may be aided by the ability to form such higher order multimers and may also serve to block access to epitopes on the more conserved S2 component of the spike, thereby facilitating immune evasion. Alternatively, loop that mediates inter-spike interactions may play a role in viral viability, consistent with our data showing that replacement of the loop with a GS linker completely abrogated viral infectivity.

We also observed two non-spike densities within the spike trimer that corresponded with linoleic acid and polysorbate 80 detergent. Linoleic acid, an essential free fatty acid, was buried within a hydrophobic pocket in the CTD with its main chain carboxyl group making contacts with the adjacent RBD in closed conformation. A recent report by Toelzer et al. also identified this density and attributed it to the presence of linoleic acid (*34*). The second large density occupied by PS 80 is situated in the NTD and is relatively more surface exposed. Since PS 80 is unique to the formulation of the Novavax 3Q-2P-FL immunogen, this observation is specific to this structure. However, there is a possibility of other ligands occupying this pocket in the place of PS 80. The presence of these binding pockets for different ligands in the spike structure provide potential targets for drug design against SARS-CoV-2.

The widely used SARS-CoV-2 spike ectodomain construct with mutated cleavage site and 2P substitution has been shown to partially exist in all RBD ‘down’ conformation or in one RBD ‘up’ conformation (*17, 18*). Surprisingly, we observed that all the RBDs in the 3Q-2P-FL spike immunogen were present in a down confirmation, which could be a cause for concern for eliciting neutralizing antibodies that compete with ACE2 binding. However, binding analysis of the 3Q-2P-FL immunogen to ACE2 by both bio-layer interferometry and ELISA clearly show binding to ACE2, indicating that the RBD is dynamic and the receptor binding site accessible (*31*). Another study on the prefusion structure of a full-length spike protein reported similar findings with RBDs clamped down as a consequence of potential clashes between S2 residues 828-853 and SD1 when RBD is in open conformation (*35*). It is also possible that the interprotomeric contacts made by linoleic acid observed in our structure preferentially lead to the observed RBD positioning. Toelzer et al. also saw a preference for their structure with linoleic acid to be in the ‘down’ state (*34*). Recent reports by Henderson et al. have revealed that mutations at certain positions can alter the propensity toward ‘up’ and ‘down’ states of the RBD (*36, 37*). They showed that introducing a disulphide bond between RBD and S2 at positions S383 and D985 stabilized the spike in an all RBD-down conformation while introducing other mutations in SD1 and S2 could alter the number of RBDs present in the ‘up’ conformation (*36*). Alternatively, the removal of N-linked glycosylation at positions 165 and 234 also resulted in an increase or decrease in the populations of spikes in the ‘up’ state, respectively (*37*).

Our structural work is consistent with the burgeoning body of structures available of the spike protein, albeit with the important differences described above. Hence, this advanced protein subunit vaccine candidate currently being tested in humans appears stable, homogeneous, and locked in the antigenically preferred prefusion conformation. Further, the tight clustering of the spikes in the nanoparticle formulation may lead to stronger immune responses over soluble trimers alone, consistent with other viral glycoprotein immunogens (HA and RSV F) (*38, 39*). It appears that the remarkable speed at which this vaccine was designed did not compromise the quality of the immunogen, and that building off the previous success in formulating RSV F and influenza HA nanoparticle immunogens could readily be extended to SARS-CoV-2 spike, particularly in the background of the 2P mutations previously shown to stabilize many other β-CoV spikes (*40*). With structural, biophysical, and antigenic characterization now complete, evaluation in humans will provide the true proof-of-principle for this vaccine concept.

## ACKNOWLEDGMENTS

We thank Bill Anderson, Hannah L. Turner and Charles A. Bowman for their help with electron microscopy, data acquisition and data processing. We thank Bill Webb and Linh Truc Hoang for their assistance with mass spectrometry and data processing. We thank Lauren Holden for her assistance with the manuscript. Authors would also like to thank Ann M. Greene at Novavax, Inc. for editing the manuscript.

## Author contributions

S.B and A.B.W conceived and designed the study. S.B, H.L.T, G.O and A.A performed cryo-EM data collection, data processing and model building. X.W, J.D, J.R.Y III and J.C.P performed site-specific glycan analysis and data interpretation. J.L.T, D.H and D.N performed mutagenesis and pseudovirus assays. S.B, G.O and A.B.W analyzed and interpreted data. S.B and A.B.W wrote the paper and all authors reviewed and edited the paper. J.H.T, A.D.P, N.P, M.J.M, G.G, and G.S contributed NVX-CoV2373 and Matrix-M adjuvant and provided advice for sample handling. J.H.T, A.D.P, N.P, M.J.M, G.G, and G.S also contributed to drafting of the manuscript.

## Data and materials availability

The EM maps have been deposited at the Electron Microscopy Data Bank (EMDB) with accession codes EMD-22352 (SARS-CoV-2 3Q-2P-FL spike trimer with C3 symmetry), EMD-22353 (SARS-CoV-2 3Q-2P-FL spike trimer with C1 symmetry), EMD-22354 (SARS-CoV-2 3Q-2P-FL spike dimer-of-trimers with C2 symmetry), EMD-22355 (SARS-CoV-2 3Q-2P-FL spike trimer-of-trimers with C1 symmetry) and EMD-22356 (SARS-CoV-2 3Q-FL spike trimer with C3 symmetry). The atomic models have been deposited at the Protein Data Bank with PDB IDs 7JJI (SARS-CoV-2 3Q-2P-FL spike trimer with C3 symmetry) and 7JJJ (SARS-CoV-2 3Q-2P-FL spike dimer-of-trimers with C2 symmetry).

## SUPPLEMENTARY MATERIALS

### Materials and Methods

#### Recombinant full-length SARS-CoV-2 prefusion S

SARS-CoV-2 full-length constructs were synthesized from the S glycoprotein gene sequence (GenBank MN908947 nucleotides 21563-25384). The wild-type full-length gene was codon optimized for expression in *Spodoptera frugiperda* (Sf9) cells by GenScript (Piscataway, NJ, USA). Amino acid mutations in the S1/S2 cleavage domain were introduced in the furin cleavage site (RRAR to QQAQ) to be protease resistant (SARS-CoV-2 3Q S) and in a second construct two proline mutations (K986P and V987P) were introduced in the HR1 domain (SARS-CoV-2 3Q-2P S) to stabilize the envelope proteins in a prefusion conformation (*18*).

#### Expression and purification

. For expression of SARS-CoV-2 full-length spike (S) proteins, the synthetic S-genes were codon optimized for insect cells, cloned into the pBac1 baculovirus transfer vector (Millipore, Sigma), and co-transfected into *S. frugiperda* [Lepidoptera] Sf9 cells with the flashBACTM GOLD system (Oxford Expression Technologies) using X-tremeGENE HP transfection reagent (Roche). Recombinant baculovirus-infected cells were harvested by centrifugation, SARS-CoV-2 S envelope proteins extracted with non-ionic detergent and purified using anion exchange and lentil lectin affinity column chromatography. Purified SARS-CoV-2 3Q and 3Q-2P S proteins were dialyzed in 25 mM sodium phosphate (pH 7.2), 300 mM NaCl, 0.01% (v/v) polysorbate 80 (PS 80) and stored at −80 °C.

#### Ns-EM sample preparation and data collection

Equal concentrations of SARS-CoV-2-3Q-2P full-length spike formulated in PS80 and Matrix adjuvant were diluted to approximately 20 µg/mL with TBS. The sample was directly deposited onto carbon-coated 400-mesh copper grids and stained immediately with 2% (w/v) uranyl formate for 90 seconds. Grids were imaged at 120 KeV on Tecnai T12 Spirit with a 4k x 4k Eagle CCD camera at 52,000x magnification and −1.5 µm nominal defocus. Micrographs were collected using Leginon and the images were transferred to Appion for processing (*41, 42*). Particle stacks were generated in Appion with particles picked using a difference-of-Gaussians picker (DoG-picker) and 2D classes generated by MSA/MRA (*43, 44*).

#### Cryo-EM sample preparation

For SARS-CoV-2 3Q-2P, 3.5 µL of spike at 0.4 mg/mL was mixed with 0.5 µL of 0.04 mM lauryl maltose neopentyl glycol (LMNG) solution immediately before sample deposition onto a 1.2/1.3 300-Gold grid (EMS) that had been plasma cleaned for 7 seconds using a Gatan Solarus 950 Plasma system. Following sample application, grids were blotted for 3 seconds before being vitrified in liquid ethane using a Vitrobot Mark IV (Thermo Fisher). For SARS-CoV-2 3Q, 3.5 µL of spike at 0.5 mg/mL was mixed with 0.5 µL of 0.04 mM of LMNG and frozen in a similar manner.

#### Cryo-EM data collection and processing

For collection, a Talos Arctica TEM at 200 kV was used with a Gatan K2 Summit detector at a magnification of 36,000x, resulting in a 1.15 Å pixel size. Total exposure was split into 250 ms frames with a total cumulative dose of ~50 e_-_/Å^2^. Micrographs were collected through Leginon software at a nominal defocus range of −0.5 µm to −1.6 µm for 3Q-2P spike and at a defocus range of −0.4 µm to −2.3 µm for 3Q spike (*45*). MotionCor2 was used for alignment and dose weighting of the frames (*46*). Micrographs were transferred to CryoSPARC 2.9 for further processing (*47*). CTF estimations were performed using GCTF and micrographs were selected using the Curate Exposures tool in CryoSPARC based on their CTF resolution estimates (cutoff 5 Å) for downstream particle picking, extraction and iterative rounds of 2D classification and selection (*48*). Particles selected from 2D classes were used for 3D refinement of free trimers for 3Q-2P-FL and 3Q-FL datasets in CryoSPARC. Final subsets of clean trimer particles were refined with C3 symmetry and local resolution for the free trimer was calculated using the local resolution function in CryoSPARC. Particles corresponding to dimers-of-trimers classes in CryoSPARC were transferred to Relion 3.0 for iterative rounds of 3D classification to separate dimers-of-trimers and trimers-of-trimers (*49*). Final subsets of clean particles from dimers-of-trimers class were refined with C2 symmetry and the trimers-of-trimers class with C1 symmetry.

#### Model building and refinement

The 3.6 Å C3-symmetric free trimer map and the 4.5 Å C2-symmetric dimers-of-trimers maps were used for model building and refinement. Initial model building was performed manually in Coot using PDB 6VXX as a template followed by iterative rounds of Rosetta relaxed refinement and Coot manual refinement to generate the final models (*50, 51*). EMRinger and MolProbity were run following each round of Rosetta refinement to evaluate and choose the best refined models (*52, 53*). To prepare linoleic acid and PS 80 detergent ligands for modeling, PDB and CIF ligand definition files were created using Phenix eLBOW (*54*) by providing the SMILES string for PubChem CID: 5284448 (polysorbate 80) or the PDB chemical component code EIC (linoleic acid). The coordinates were manually placed and refined into the respective map densities using Coot. For Rosetta refinement, each ligand was saved in MOL2 format and Rosetta parameter files were generated using the molfile_to_params.py function (*51*). Final map and model statistics are summarized in Table S1. Figures were generated using UCSF Chimera and UCSF Chimera X (*55, 56*).

#### Mass-spectrometry

Mass spectrometry to identify fatty acids in the SARS-3Q-2P-FL protein was performed as described previously (*21*). We obtained several candidates in this screen that were narrowed down to 6 candidates based on their intensity and the m/z range of 250-300.

#### Sequence alignment

The sequences for the analysis were obtained either from GISAID or NCBI GenBank. GISAID accession number or the GenBank accession id from which whole genome sequences were obtained are as follows: Bat-SL-RatG13 (EPI_ISL_402131), Bat-SL-CoVZC45 (MG772933.1), BetaCoV/pangolin/Guandong/1/2019 (EPI_ISL_410721) and BetaCoV/pangolin/Guangxi/P4L/2007 (EPI_ISL_410538), SARS-CoV Tor2 strain (NC_004718.3). The spike sequences were extracted and aligned using Clustal Omega (*57*).

#### Pseudovirus (PSV) assay

Pseudovirus preparation and assay were performed as previously described (*58*). Under BSL2/3 conditions, MLV-gag/pol and MLV-CMV plasmids was co-transfected into HEK293T cells along with full-length or mutant SARS-CoV-2 spike plasmids using Lipofectamine 2000 to produce a single round of infection competent pseudo-viruses. The medium was changed 12 hours after transfection. Supernatants containing the viruses were harvested 48h after transfection. In sterile 96-well half-area plates, 25 µl of virus was immediately added to 10,000 HeLa or HeLa-ACE2 cells in 75 µl of medium. Plates were incubated at 37°C for 42 to 48 h. Following the infection, HeLa and HeLa-hACE2 cells were lysed using 1x luciferase lysis buffer (25 mM Gly-Gly pH 7.8, 15 mM MgSO_4_, 4 mM EGTA, 1% Triton X-100). Luciferase intensity was then read on a Luminometer with luciferase substrate according to the manufacturer’s instructions (Promega, PR-E2620).

#### Site-specific glycosylation

A sample of the SARS-CoV-2 prefusion spike protein expressed in the SF9 insect cell line was prepared for MS analysis as previously described with minor modifications (*24*). In brief, the protein (50 µg) was denatured and aliquots (10 µg) were digested under five different protease conditions including chymotrypsin, a combination of trypsin and chymotrypsin, trypsin, elastase and subtilisin as described. All samples were then pooled and deglycosylated by Endo H followed by PNGase F in O_18_-water. To obtain full site coverage, an additional aliquot of the denatured protein (10 µg) sample was digested with chymotrypsin (1:13 w/w) only and deglycosylated with EndoH and PNGase F like the other samples.

The combined protease-treated and chymotrypsin only samples were separately analyzed on an Q Exactive HF-X mass spectrometer (Thermo). Each sample was run twice as replicate. Samples were injected directly onto a 25 cm, 100 µm ID column packed with BEH 1.7 µm C18 resin (Waters). Samples were separated at a flow rate of 300 nL/min on a nLC 1200 (Thermo). Solutions A and B were 0.1% formic acid in 5% and 80% acetonitrile, respectively. A gradient of 1–25% B over 160 min, an increase to 40% B over 40 min, an increase to 90% B over another 10 min and held at 90% B for 30 min was used for a 240 min total run time. Column was re-equilibrated with solution A prior to the injection of sample. Peptides were eluted directly from the tip of the column and nanosprayed directly into the mass spectrometer by application of 2.8 kV voltage at the back of the column. The HFX was operated in a data dependent mode. Full MS1 scans were collected in the Orbitrap at 120k resolution. The ten most abundant ions per scan were selected for HCD MS/MS at 25NCE. Dynamic exclusion was enabled with exclusion duration of 10 s and singly charged ions were excluded.

The MS data were processed essentially as described previously (*24*). The data were searched against the proteome database and quantified using peak area in Integrated Proteomics Pipeline-IP2. Since the processing pathway in SF9 cell line (insect cell line) is similar to mammalian cells for oligomannose and hybrid structures cleaved by Endo-H, and then diverges to produce a combination of paucimannose and complex type glycans, peptides with N+203 were identified as having oligomannose type glycans, and peptides with N+3 are assigned as peptides with complex and paucimannose type glycans.

**Figure S1.**
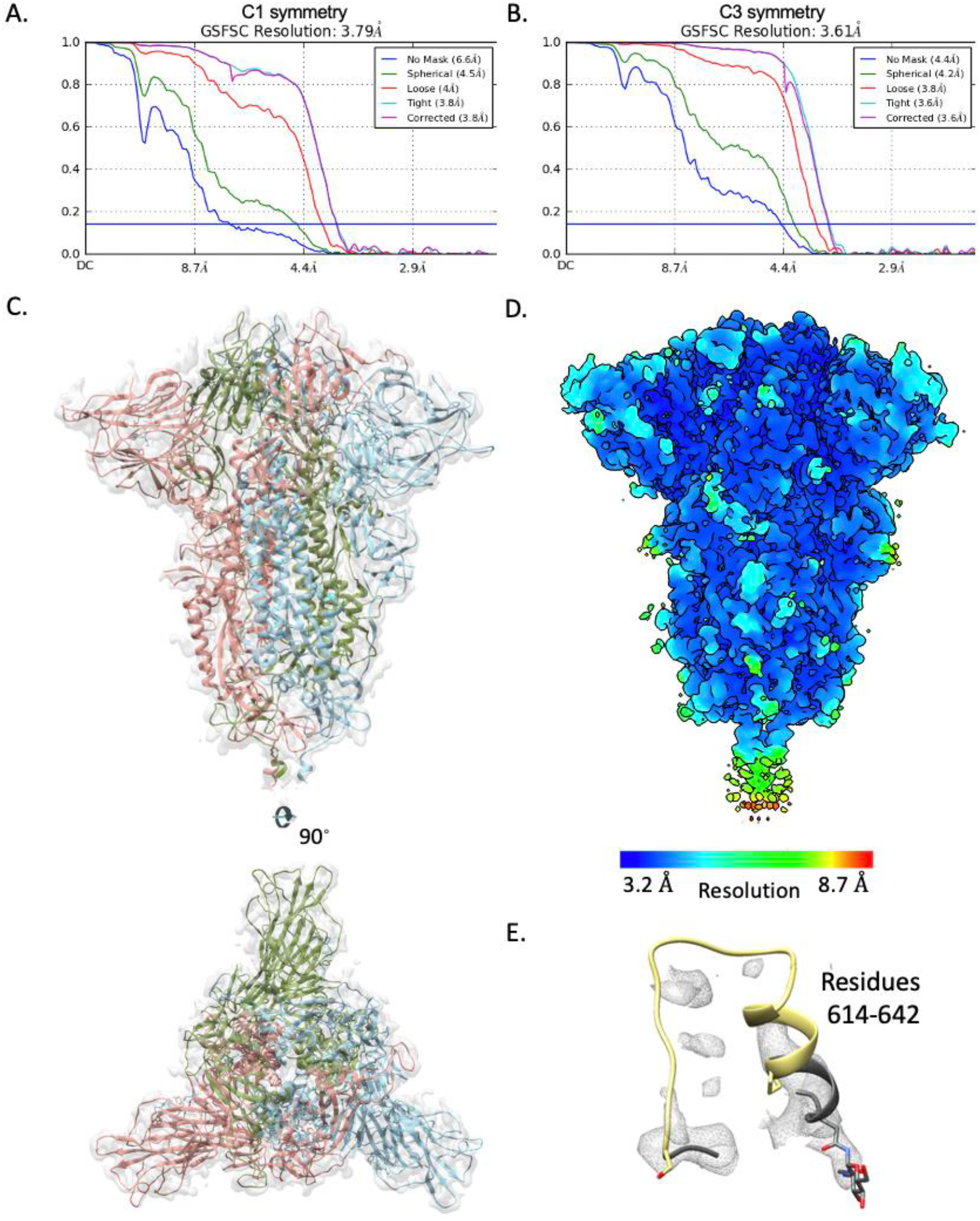
Cryo-EM structure analysis and validation of SARS-CoV-2 3Q-2P-FL spike trimer. **(A)** FSC curve for SARS-CoV-2 3Q-2P-FL spike when C1 symmetry was imposed during refinement. **(B)** FSC curve for SARS-CoV-2 3Q-2P-FL spike with C3 symmetry imposed. **(C)**Side and top view of the C3 trimer atomic model represented as a ribbon diagram fit into the 3Q-2P-FL spike C1 map density. The protomers are colored in blue, green and coral and the map is shown as a transparent gray density. **(D)** Cryo-EM map of SARS-CoV-2 3Q-2P-FL C3, colored according to local resolution. **(E)** Ribbon representation of residues 614-642 of the PDB model 6X6P with their corresponding cryo-EM density (EMD-22078) shown as mesh representation. Residues P621 to S640 are colored in yellow.

**Figure S2.**
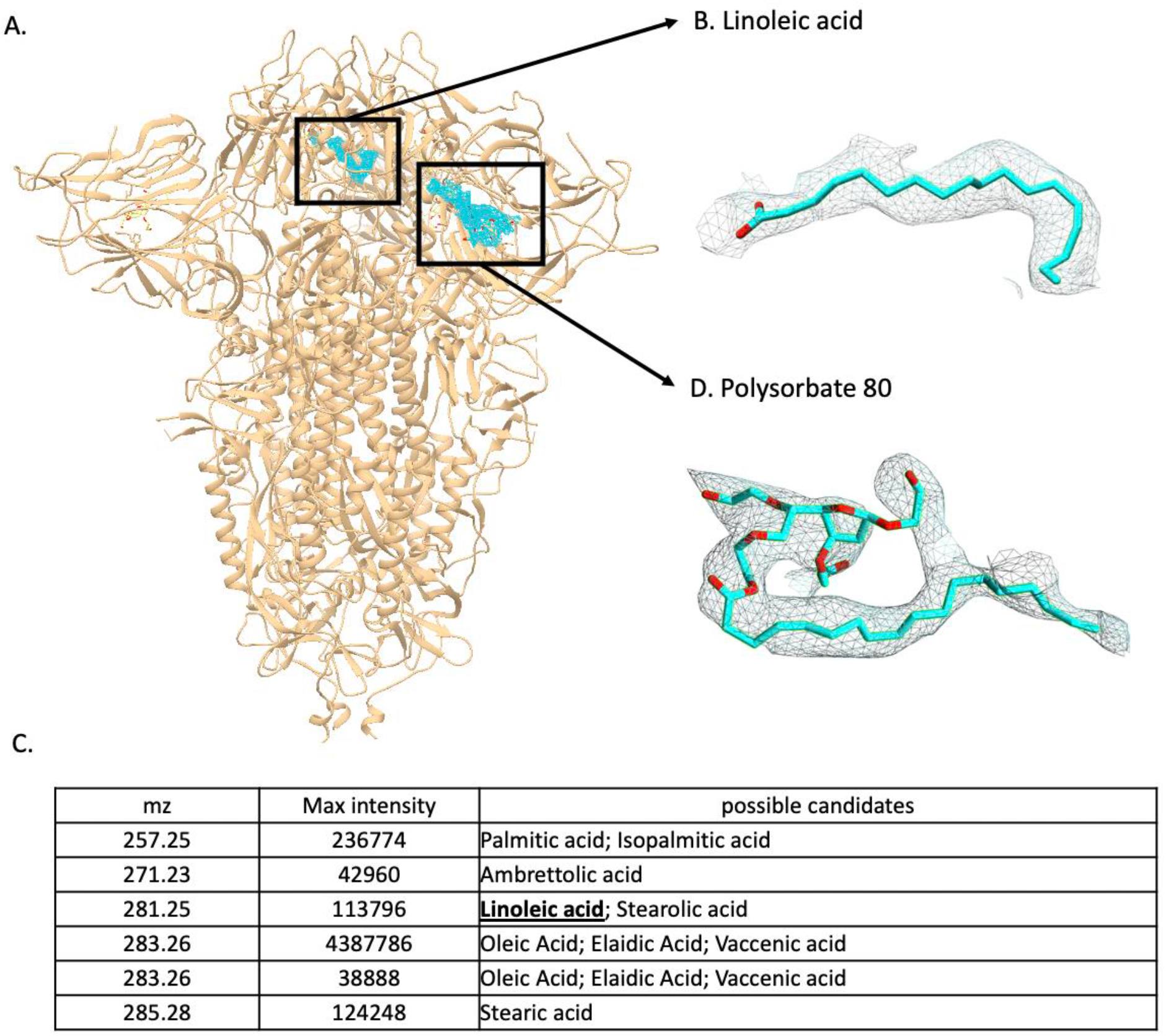
Ligands found in the EM map of the SARS-CoV-2 3Q-2P-FL spike trimer. **(A)**Side view of the trimer atomic model represented as a ribbon diagram in tan with the two ligands and their corresponding map densities shown as mesh in cyan. **(B)** Linoleic acid and its corresponding map density shown as mesh in cyan. **(C)** The potential ligand candidates obtained from the mass spectrometry analysis of the SARS-CoV-2 3Q-2P-FL spike trimer with their corresponding size and signal intensity. **(D)** Polysorbate 80 and its corresponding map density shown as mesh in cyan.

**Figure S3.**
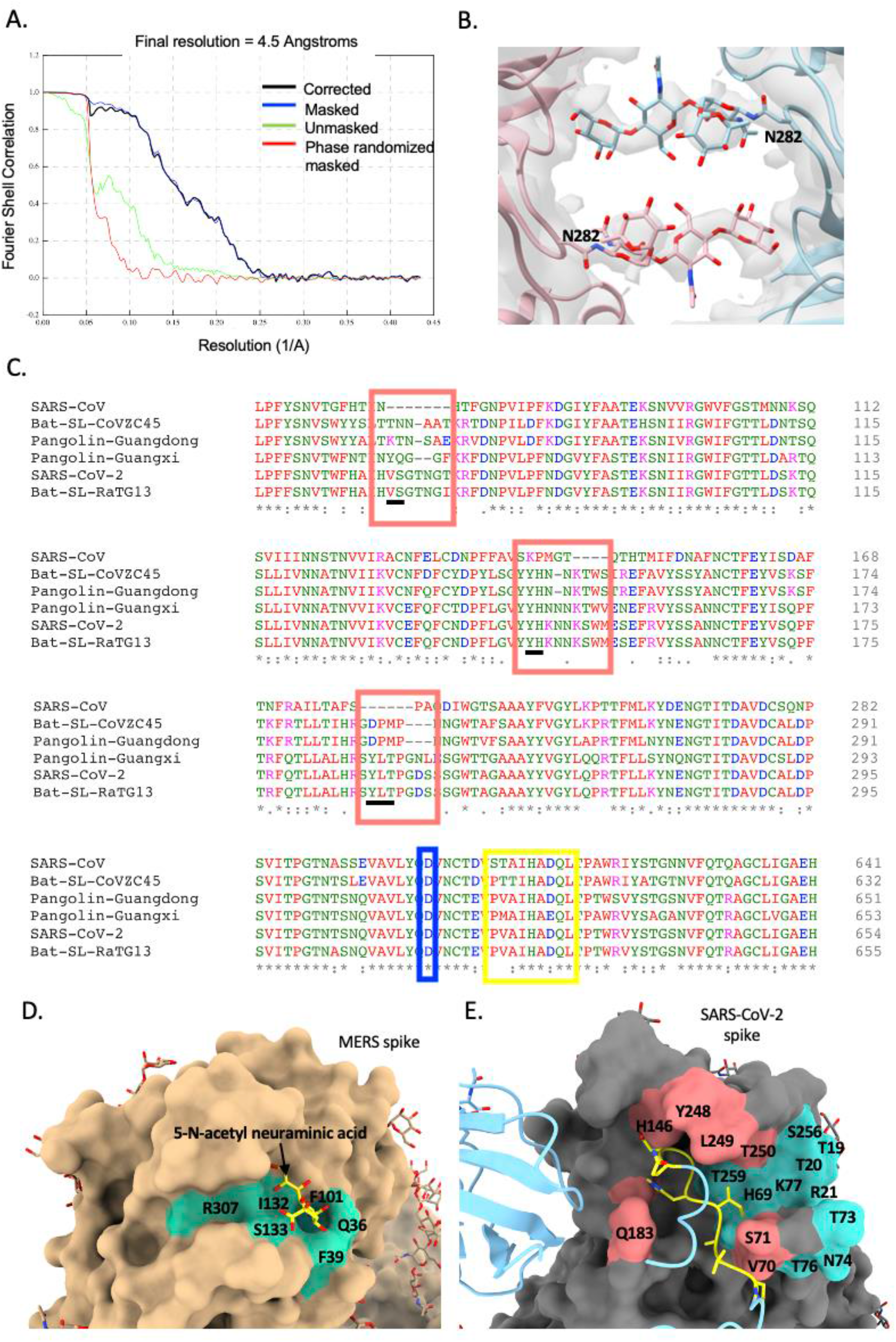
Cryo-EM validation and analysis of SARS-CoV-2 3Q-2P-FL spike dimers-of-trimers. **(A)** FSC curve for 3Q-2P-FL dimers-of-trimers map imposing C2 symmetry. N282 glycans extending out from each trimer towards the symmetry related chain in the adjacent trimer. The adjacent spike trimers are shown in pink and blue as ribbon representation and their corresponding cryo-EM density shown in transparent gray as surface representation. **(C)** Alignment of spike sequences from representative lineage B beta-CoV strains performed using Clustal Omega. The loop residues 621-PVAIHADQ-628 are highlighted by a yellow box, the D614 residue highlighted by a blue box, the loops surrounding the NTD binding pocket are highlighted by a coral box and the potential interacting residues are underlined in black. **(D)** Surface representation of MERS spike (PDB ID: 6Q04) in tan color bound to 5-N-acetyl neuraminic acid shown in yellow. The binding site is colored in cyan. **(E)** Interaction between the protomers of adjacent trimers in the 3Q-2P-FL dimers-of-trimers model. One protomer is shown as a ribbon diagram in blue while its binding partner is shown as surface representation in gray. Residues 621-PVAIHADQ-628 on the loop with potential interactions are colored yellow and the corresponding residues in the NTD binding pocket are highlighted in coral. Spike residues predicted in glycan binding are colored in cyan.

**Figure S4.**
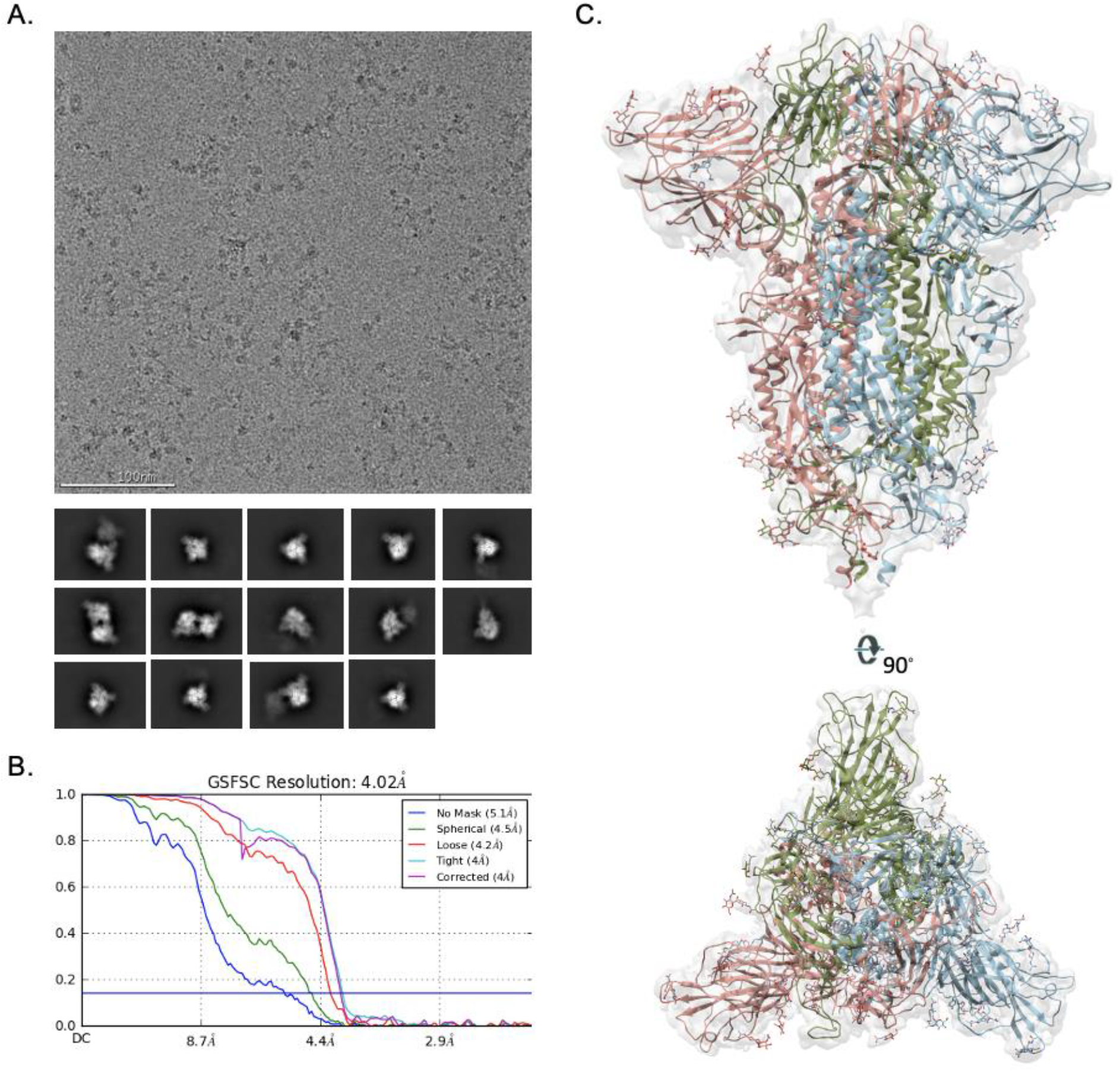
Cryo-EM validation and analysis of SARS-CoV-2 3Q-FL spike. **(A)**Representative electron micrograph and 2D class averages of 3Q-FL spikes showing free trimers and complexes of trimers. **(B)** FSC curve for SARS-CoV-2 3Q-FL spike (C3 symmetry imposed during map refinement). **(C)** Side and top view of the SARS-CoV-2 3Q-2P-FL C3 trimer atomic model represented as a ribbon diagram fit into the 3Q-FL spike C3 map density. The protomers are colored in blue, green and coral and the map is shown as a transparent gray density.

**Table S1.** Cryo-EM data collection, refinement and model building statistics.

| Map | 3Q-2P-FL C3<br>(trimer) | 3Q-2P-FL C2<br>(dimer of<br>trimers) | 3Q-2P-FL C1<br>(trimer of<br>trimer) | 3Q-2P-FL C1<br>(trimer) | 3Q-FL C3<br>(trimer) |
| --- | --- | --- | --- | --- | --- |
| EMDB | EMD-22352 | EMD-22354 | EMD-22355 | EMD-22353 | EMD-22356 |
| **Data collection** |  |  |  |  |  |
| Microscope | FEI Talos<br>Arctica | FEI Talos<br>Arctica | FEI Talos<br>Arctica | FEI Talos<br>Arctica | FEI Talos<br>Arctica |
| Voltage (kV) | 200 | 200 | 200 | 200 | 200 |
| Detector | Gatan K2<br>Summit | Gatan K2<br>Summit | Gatan K2<br>Summit | Gatan K2<br>Summit | Gatan K2<br>Summit |
| Recording mode | Counting | Counting | Counting | Counting | Counting |
| Nominal magnification | 36,000 | 36,000 | 36,000 | 36,000 | 36,000 |
| Movie micrograph pixel size (Å) | 1.15 | 1.15 | 1.15 | 1.15 | 1.15 |
| Dose rate (e-/[(camera pixel)*s]) | 5.7 | 5.7 | 5.7 | 5.7 | 5.95 |
| Number of frames per movie<br>micrograph | 46 | 46 | 46 | 46 | 44 |
| Frame exposure time (ms) | 250 | 250 | 250 | 250 | 250 |
| Movie micrograph exposure time<br>(s) | 11.5 | 11.5 | 11.5 | 11.5 | 11.1 |
| Total dose (e-/Å<sup>2</sup>) | 50 | 50 | 50 | 50 | 50 |
| Defocus range (µm) | -0.5 to -1.6 | -0.5 to -1.6 | -0.5 to -1.6 | -0.5 to -1.6 | -0.4 to -2.3 |
| **EM data processing** |  |  |  |  |  |
| Number of movie micrographs | 5,506 | 5,506 | 5,506 | 5,506 | 3,627 |
| Number of molecular projection<br>images in map | 45,374 | 15,411 | 4,141 | 45,374 | 76,834 |
| Symmetry | C3 | C2 | C1 | C1 | C3 |
| Map resolution (FSC 0.143; Å) | 3.6 | 4.5 | 8.0 | 3.8 | 4.0 |
| Map sharpening B-factor (Å<sup>2</sup>) | -114.8 | -66.5 | -240.3 | -88.8 | -123.5 |
| **Structure Building and**<br>**Validation** |  |  |  |  |  |
| *Number of atoms in deposited*<br>*model* |  |  |  |  |  |
| SARS-CoV-2 S protein | 25,971 | 52,088 | N/A | N/A | N/A |
| Glycans | 1,302 | 2,526 | N/A | N/A | N/A |
| Other ligands | 186 | 0 | N/A | N/A | N/A |
| MolProbity score | 0.76 | 0.75 | N/A | N/A | N/A |
| Clashscore | 0.48 | 0.76 | N/A | N/A | N/A |
| Map correlation coefficient | 0.85 | 0.80 | N/A | N/A | N/A |
| EMRinger score | 3.69 | 1.64 | N/A | N/A | N/A |
| *RMSD from ideal* |  |  |  |  |  |
| Bond length (Å) | 0.02 | 0.02 | N/A | N/A | N/A |
| Bond angles (°) | 1.74 | 1.79 | N/A | N/A | N/A |
| *Ramachandran plot* |  |  |  |  |  |
| Favored (%) | 98.37 | 97.95 | N/A | N/A | N/A |
| Allowed (%) | 1.63 | 2.05 | N/A | N/A | N/A |
| Outliers (%) | 0.00 | 0.00 | N/A | N/A | N/A |
| Side chain rotamer outliers (%) | 0.21 | 0.07 | N/A | N/A | N/A |
| PDB | 7JJI | 7JJJ | N/A | N/A | N/A |

